# mAFiA: Detecting m^6^A at single-molecular resolution via direct-RNA sequencing

**DOI:** 10.1101/2023.07.28.550944

**Authors:** Adrian Chan, Isabel S. Naarmann-de Vries, Carolin P. M. Scheitl, Claudia Höbartner, Christoph Dieterich

## Abstract

Direct-RNA sequencing offers the possibility to simultaneously identify canonical bases and epi-transcriptomic modifications in each single RNA molecule. Thus far, the development of computational methods has been hampered by the lack of biologically realistic training data that carries modification labels at molecular resolution. Here, we report on the synthesis of such samples and the development of a bespoke algorithm that accurately detects single m^6^A nucleotides on single molecules in both synthetic RNAs and natural mRNA.

---

Each time-trace in **d**irect-**RNA** sequencing (dRNA-Seq) on the Oxford Nanopore platform encodes the unique modification fingerprint of an individual molecule^1^. However, no database of naturalistic RNA with modification labels at the read-level exists yet. For m^6^A specifically, methods development thus far is based on either synthetic molecules with all-or-none modification patterns^2^, or biological mRNA whose modification sites are only known at the ensemble level^3^. In this work, we employ an alternative strategy by synthesizing short RNA oligos that portray actual mRNA sections with m^6^A sites, while the modification status of each nucleotide (nt) is precisely controlled.

Our synthetic samples include the six most common DRACH motifs, which collectively account for almost 80% of all consensus m^6^A sites in human mRNA^4–6^. For each motif, non-intersecting sequence designs are chosen from sections of the human transcriptome with known m^6^ A loci, in the form of 21-or 33-mers (see Table 1). Each sequence is synthesized in two variants that contain either an A or an m^6^ A in the center. To produce polymers suitable for dRNA-Seq, the oligos are ligated into longer RNA molecules using two different strategies (Figure 1a). Hereafter, we denote the random-ligated samples by “RL” and the splint-ligated ones by “SPL”.

**Figure 1:**
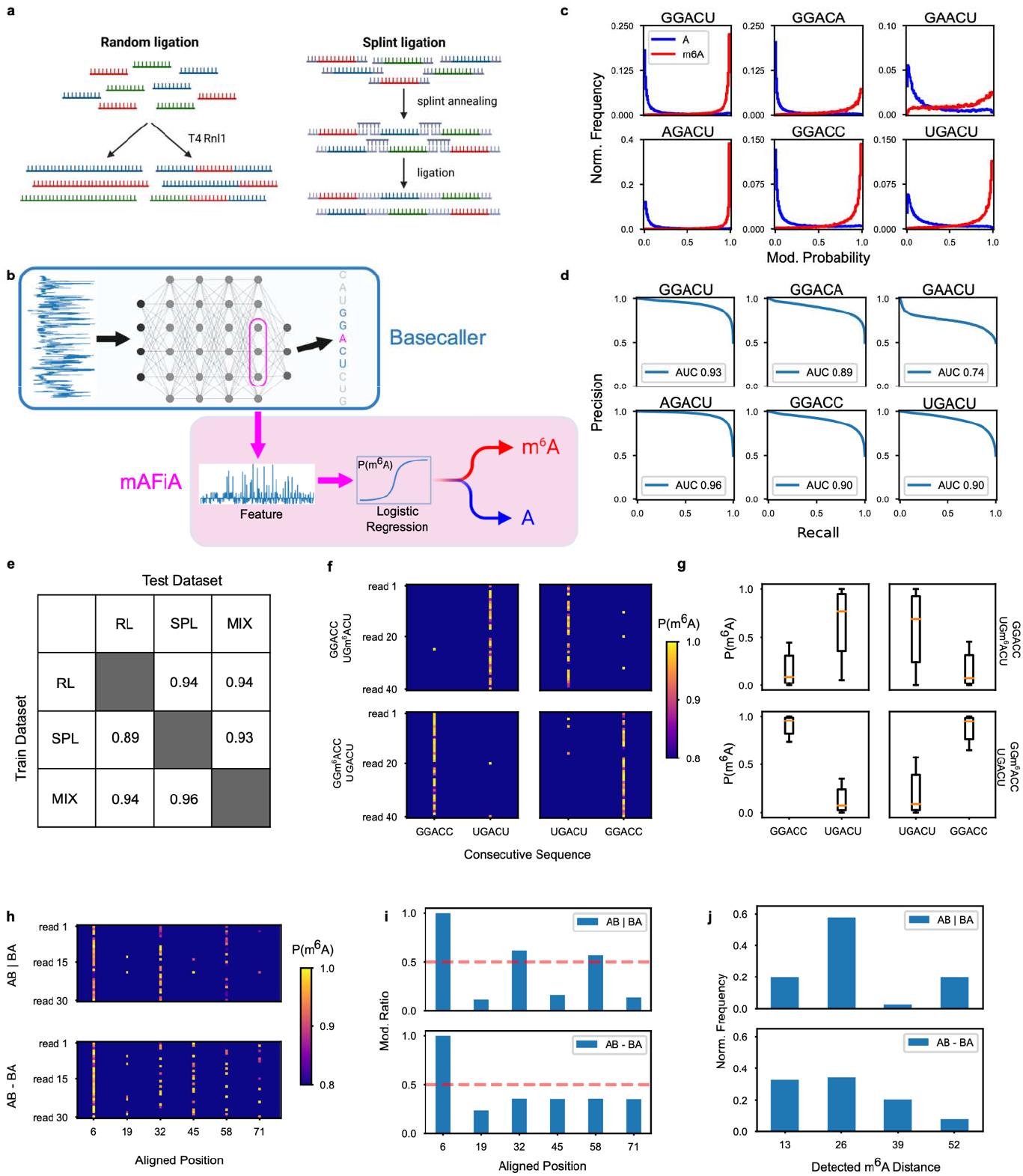
Methods and single-molecule experiments. **a**, The RL samples are concatenated by random ligation to either homo- or heteropolymers (left), the SPL samples by splint ligation to heteropolymers (right). **b**, mAFiA as an add-on module to a basecaller network. **c**, Normalized distribution of single-molecule P(m^6^ A) predicted by mAFiA, trained on SPL and tested on RL. **d**, Precision-Recall curve calculated from **c. e**, Average AUC of all six motifs with different train and test sets. **f**, Single-read P(m^6^ A) predicted on RL heteropolymers M_4_(A)M_5_(m^6^ A) and M_4_(m^6^ A)M_5_(A), showing read sections aligned to the consecutive patterns M_4_→M_5_ or M_5_→M_4_. **g**, Box plot of aggregate P(m^6^ A) of aligned locations from RL M_4_(A)M_5_(m^6^ A) and M_4_(m^6^ A)M_5_(A). **h**, Single-read P(m^6^ A) predicted on SPL AB|BA and AB-BA. Reads aligned by their first detected m^6^ A. **i**, Aggregate modification ratio in the center of each 13-nt cycle. **j**, Probability of detected m^6^ A distance on a single read.

To detect m^6^ A modification patterns in dRNA-Seq, we developed a transfer learning method with minimal computational overhead. Our **m**^6^**A Fi**nding **A**lgorithm (mAFiA) re-uses internal features generated by the backbone neural network during basecalling, and assigns an m^6^ A probability, P(m^6^ A), to a specific A on the read (Figure 1b). Unlike other single-molecule methods, mAFiA does not require additional post-processing such as nanopolish^7^, and can be integrated into an existing basecaller without altering the latter’s accuracy.

First, we cross-validated our method on the RL and SPL datasets, which involve different sequence designs and ligation strategies. Figure 1c shows the predictions of a mAFiA model trained on SPL samples and tested on RL. For each DRACH motif, there is a stark divergence in P(m^6^ A) between the modified and unmodified molecules. The precision-recall curves on 5 out of 6 motifs yield area-under-curve (AUC) values of close to or exceeding 0.9 (Figure 1d), which suggests that our model trained on one dataset performs robustly in hitherto unseen sequence contexts. We repeat the cross-validation using different permutations of train and test datasets, and evaluate the model’s performance using the average AUC of 6 motifs. Not surprisingly, the model resulting from a joint dataset performs better than those trained on either sample alone (Figure 1e). Thereafter, all evaluations are performed using the combined model.

To ascertain that our method can indeed locate m^6^ A at single-molecule, single-nucleotide resolution, we designed two further experiments in which the RNAs are indistinguishable in their base sequences, but differentiable only in their underlying modification patterns. The first experiment involves heteropolymers containing RL oligos M_4_ (GGACC) and M_5_ (UGACU) in equal proportions, with m^6^ A inserted in either one of the two motifs. Figure 1f shows the read-by-read predicted P(m^6^ A) in each sample, showing read sections mapped to the consecutive sequence pattern M_4_ immediately followed by M_5_, or M_5_ followed by M_4_. For each combination of insertion location and alignment pattern, one can observe a distinct high P(m^6^ A) location on most reads that corresponds to the modified location. Crucially, for the same aligned sequence, there is a clear reversal of the P(m^6^ A) pattern as the modified site switches side (Figure 1g). The resulting four-fold mirror symmetry leaves no doubt that the detected m^6^ A indeed reflects underlying modification signatures rather than specific sequence contexts.

The second single-molecule experiment involves two samples, AB|BA and AB-BA, which entail the motif GGACU with A or m6A inserted in its center (Figure S3a). Both samples contain periodic repetition of the same 13-mer sequences. However, AB|BA only allows m^6^ A spacing of 26 nts, while AB-BA admits all spacing in multiples of 13 nts. Figure 1h shows the read-by-read P(m^6^ A) detected in each sample, which indeed shows synchronized periods of high P(m^6^ A) in AB|BA but not in AB-BA. The picture becomes more apparent as we calculate the aggregate modification ratio at each aligned location (figure 1i). After the first detected m^6^ A, AB|BA exhibits oscillating modification ratios at a period of 26 nts, while AB-BA does not. On any single read, the characteristic m^6^ A separation in AB|BA occurs at 26 nts, while the same is not observed in AB-BA (figure 1j). To establish a benchmark for all single-molecule methods, we define a metric K (see methods) that measures the contrast in m^6^ A distance distribution. K=1 indicates perfect contrast, while K=0 indicates no detected difference between AB|BA and AB-BA. mAFiA attains a score of 0.66. Two other methods, CHEUI^2^ and m6Anet^3^, obtain best scores of 0.40 and 0.22 respectively (Figure S3b-c).

Having validated our method on synthetic molecules, we applied it to biological RNA. While per-read labels are unavailable for such data, ensemble-level stoichiometry has been estimated for HEK293 mRNA with GLORI^8^. Figure 2a shows an example of read-by-read mAFiA prediction for a pair of m^6^ A sites on the same gene. Despite their close proximity, a stark contrast in P(m^6^ A) distribution between these two sites is observed (Figure 2b). Using 0.5 as the modification threshold, we calculate the stoichiometry at each site, and compare it to the value reported by GLORI (Figure 2c). mAFiA obtains a correlation of 0.85. On the same intersecting set of sites, the predictions of CHEUI and m6Anet yield correlations of 0.64 and 0.80 respectively (Figure S3d-e). At both single-molecule and ensemble level, mAFiA outperforms existing methods (Table 4).

**Figure 2:**
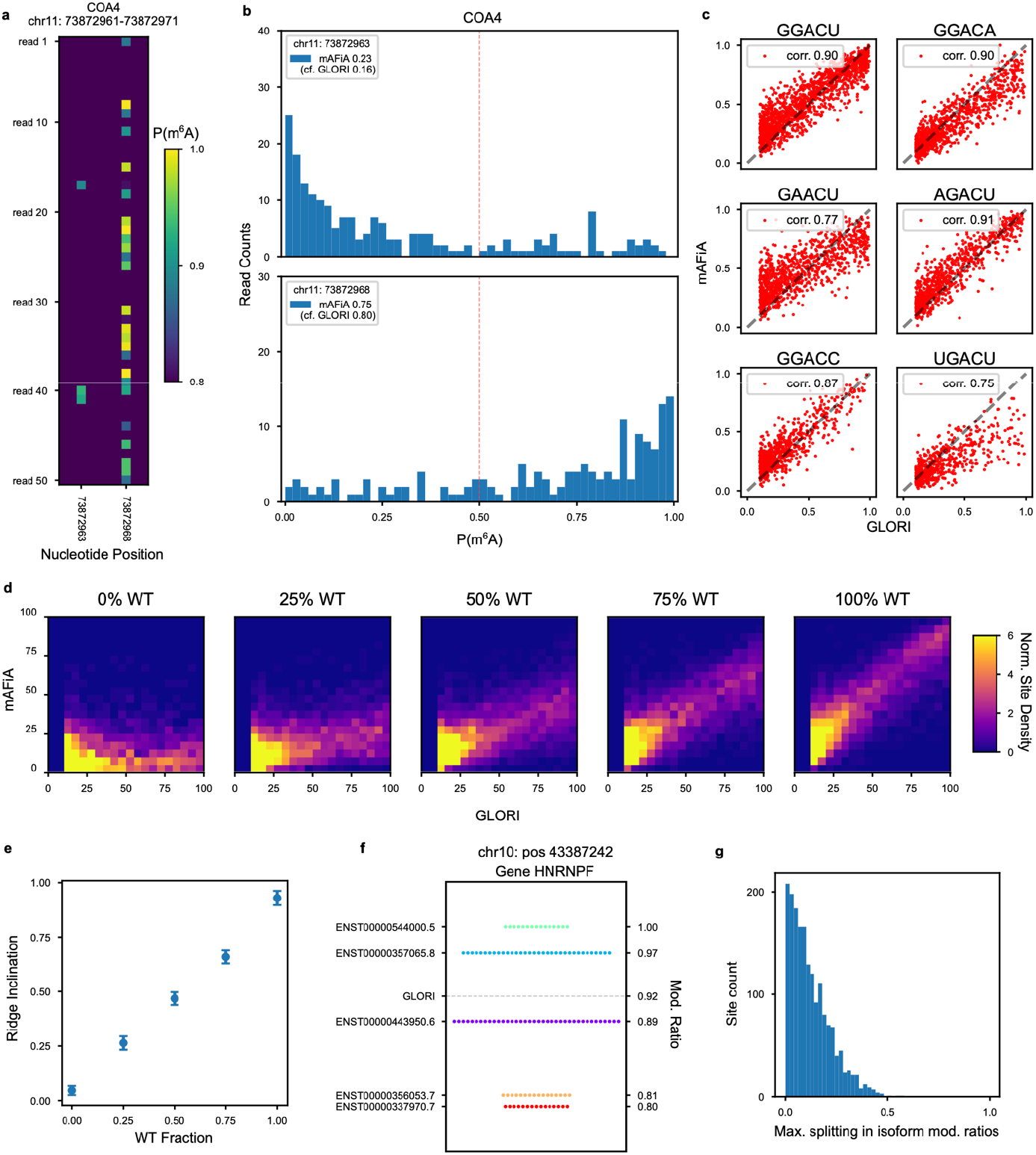
Testing on HEK293 mRNA. **a**, Single-read P(m^6^ A) predicted on a pair of proximate sites on COA4. **b**, Histogram of all P(m^6^ A) on the same pair of sites. Legend: Site-level modification ratio calculated with threshold 0.5. GLORI reference value in brackets. **c**, Site-level stoichiometry comparison between mAFiA and GLORI for six DRACH motifs. **d**, 2D-density plots of site-level mAFiA-GLORI comparison at various concentrations of WT-IVT mixing. **e**, Slope of ridge in **d** versus WT percentage. **f**, m^6^ A stoichiometry predicted by mAFiA for variou transcripts aligned to the same genomic location on HNRNPF. Each dot represents one read. GLORI (gray dashed line) represents a weighted average between various isoforms. **g**, Distribution of maximum difference in isoform stoichiometry within each measured genomic site.

In addition to the HEK293 wildtype (WT), we also prepared mixtures of WT with *in vitro* transcribed (IVT) RNA at different ratios. As the IVT sample is devoid of RNA modifications, we expect the overall stoichiometry of each site to scale accordingly with WT-IVT ratio. The results displayed in Figure 2d resoundingly confirm our hypothesis, where the slopes fitted to respective mAFiA-GLORI correlation ridges correspond almost exactly to the percentage of WT in the mixture (Figure 2e).

An additional advantage of dRNA-Seq is its ability to distinguish between different transcript isoforms. Interestingly, for many m^6^ A sites, the reference stoichiometry reported by GLORI represents merely a weighted average between various splice variants (Figure 2f). Overall, the divergence in modification ratios between isoforms at the same site can reach up to 0.5 among the sites that we have measured (Figure 2g). We repeated our measurements with deeper coverage, and observed that the isoform-specific stoichiometry predicted by mAFiA is highly reproducible (Figure S4).

Through meticulous validation in both synthetic and natural RNAs, we have demonstrated an accurate m^6^ A finding algorithm at single-molecule resolution. We hope this will be the first step towards a multi-modification detector as new data comes online^9^. Further, our method opens up the opportunity for isoform-specific stoichiometry, which we will continue to investigate with follow-up experiments.

