## Supplementary material for "mAFiA: Detecting m^6^A at single-molecular resolution via direct-RNA sequencing": Methods and Supplement

#### RNA oligos

Synthetic RNA oligos with a 5' phosphate were purchased or prepared in house by solid-phase synthesis using 2'-*O*-TOM-protected RNA phopshoramidite as previously described[1]. Oligos for random ligation (GenScript, purification grade) were of 21 nts length; oligos for splinted ligation were 33 nts in length. Each oligo was synthesized either as unmodified oligo or with an $m^6A$ at the central position. The oligos cover the six most abundant DRACH motifs according to GLORI-seq. The 5' and 3' context was chosen based on detected frequency and ligation efficiency. Oligos were dissolved at 100 µM in nuclease-free water and stored at -80°C (Table 1).

#### Optimization of Random ligation

The reaction conditions for random ligation were optimized to increase the fraction of linear products and decrease the circular RNA formation. Starting from the manufacturers protocol (https://international.neb.com/protocols/2018/10/17/protocol-ligation-of-an-oligo-to-the-3-end-of-rna-using-t4-rna-ligase-1m0204), a) the reaction temperature was decreased from 25°C to 4°C, b) the reaction time was increased from 2 h to 6h + 16 h c) the ATP concentration was decreased from 1 mM to 0.1 mM, d) no benefit for higher PEG8000 concentration than 25% nor DMSO addition was identified and e) the oligo amount was increased from 40 pmol to 1000 pmol. The improvement in random ligation is shown in Figure S1.

#### RNase R treatment

The relative proportion of circular product was determined by RNase R digestion, which degrades all linear RNA. 10 µl ligation reaction was digested with 1 µl RNase R (Lucigen) in a total volume of 20 µl for 30 min at 37°C. The enzyme was inactivated 20 min at 65°C. Comparable amounts of undigested and RNase R treated samples were analyzed on 10% TBE-urea gels.

#### Random Ligation and polyadenylation

First, random ligation was optimized (outlined below). 1 nmol RNA oligo with 25% PEG8000 and 0.1 mM ATP in 1x T4 RNA ligase buffer and 10 U T4 RNA ligase 1 (New England Biolabs) was ligated for 6 h on ice. After this, additional 10 U T4 RNA ligase 1 were added and ATP increased to 0.2 mM. Ligation was carried on overnight at 4°C. Ligation products were purified with RNA Clean & Concentrator-5 (Zymo Research) according to the standard protocol. Then, ligation products should be sequenced together were mixed at approximately equimolar ratio and polyadenylated with E-PAP Poly(A) Tailing Kit (Thermo Fisher Scientific) in a total volume of 50 µl with 1.5 µl RiboLock (Thermo Fisher Scientific) and 2.5 µl E-PAP for 5 min at 37°C. Subsequently, polyadenylated mixtures were purified with RNA Clean & Concentrator kit-5 according to the standard protocol. Samples were either subjected directly to library preparation or stored at -80°C until further usage.

### PAGE analysis of random ligation products

Aliquots of the ligation reaction were analyzed on 10% TBE-urea polyacrylamid gels (Novex, Thermo Fisher Scientific) to assess ligation efficiency. 2 µl ligation reaction or low range ssRNA ladder (New England Biolabs) were heated for 5 min to 70°C in 10 µl Novex TBE-urea sample buffer and rapidly cooled before loading. Additionally, 1 µl GeneRuler Ultra Low Range DNA Ladder (Thermo Fisher Scientific) was used for size estimation. Gels were stained 15 min at room temperature with SYBR Green II RNA gel stain (Thermo Fisher Scientific) and images acquired on a ChemiDoc Imaging System (Bio-Rad).

### Splinted Ligation

For concatenation by splinted ligation, 500 pmol RNA (containing equal amounts of the individual RNA sequences) and 500 pmol of the DNA splint (ordered from Microsynth, see Table 3) were annealed (5 min at 95 °C, 10 min at 25 °C) in annealing buffer (4 mM Tris-HCl (pH 8.0), 15 mM NaCl, 0.1 mM EDTA). Then, $MgCl_2$ was added to a final concentration of 8 mM along with 1.5 µL 10x T4 RNA ligase 2 buffer and 15 U of the T4 RNA ligase 2 (New England Biolabs). Ligation reactions were carried out in a final reaction volume of 15 µL. The ligation reactions were incubated overnight at 25 °C, followed by loading on 15 % denaturing PAGE (distributed into 5 wells on a 85 x 70 x 1 mm gel, 200 V, 50 min) with running buffer 1x TBE (89 mM Tris, 89 mM boric acid, 2 mM EDTA, pH 8.3). Synthetic RNAs of 35, 104 and 193 nts were used as size markers. The gels were stained with SYBR green and imaged on a ChemiDoc Imager (Bio-Rad). Ligation products > 150 nt in length were excised and extracted with TEN buffer (10 mM Tris-HCl, pH 8.0, 1 mM EDTA, 300 mM NaCl). The RNA was recovered by precipitation with cold ethanol, yielding around 20-25 ng ligated RNA product per 15 µL ligation reaction. Polyadenylation was done as for randomly ligated products described above.

### Direct RNA-seq library preparation and ONT sequencing

Direct RNA-sequencing libraries were prepared from approximately 1 µg of polyadenylated mixtures with RNA-SQK002 (Oxford Nanopore Technologies, ONT hereafter) with some modifications as described in https://www.biorxiv.org/content/10.1101/2021.11.10.467884v1. Ligation reactions were done for 10 min at room temperature. Libraries were quantified with the dsDNA HS Qubit kit (Thermo Fisher Scientific) and loaded completely on Flongle flow cells (R9.4.1). Sequencing was performed for 24 h in MinION devices connected to a Dell workstation (specifications) with high accuracy live basecalling and bulk output turned on.

### Backbone network

The backbone basecaller is based on the RODAN architecture[2] with several adaptations. First, the input length of the raw signal is shortened to 1024. Second, the decoder uses a Viterbi[3] rather than the default CTC algorithm[4]. Third, the model is trained on IVT HEK293 mRNA instead of a mixture of WT data from different species. Data preparation follows the walkthrough of the ONT software Taiyaki[5]. Training procedure follows the instructions on RODAN's repository (https://github.com/biodlab/RODAN ).

### mAFiA module

Feature vectors of length 768 are extracted from the last convolution layer ('convlayers.conv21') of the backbone network. Each vector is normalized by its maximum absolute value. A linear logistic model[6] is then fitted to the feature to produce a probability between 0 and 1.

### Training on synthetic molecules

dRNA-Seq data from both RL and SPL samples are basecalled with the backbone network. Oligo sequences are mapped locally to each read and then chained together to form a ligated sequence. For each of the six DRACH motifs, features that correspond to the locations of the central A nucleotide are collected, then split into 0.75/0.25 train/test sets. Training of the logit classifier proceeds with the default L-BFGS solver[7] and maximum iterations 1000.

### Single-molecule benchmark (AB+BA)

Basecalled reads are aligned with minimap 2.22 to reference sequences that consist of repeating patterns of 26-mer (see table 2), which range from 1 to 10 cycles (26-260 nts). The first detected $m^6A$ in each read is aligned to position 6, the center of the first GGACU motif. Reads that contain fewer than 2 detected $m^6A$'s are excluded.

The distance between a pair of detected $m^6A$'s is the number of nucleotides between them in the aligned read. The collection of all $m^6A$ distances on single reads are then aggregated into a frequency distribution f(x), where x can be 13, 26, 39, or 52 nts. f(x) is normalized such that its sum over x is equal to 1. The contrast k on a sample is defined as:

$$k = f(26) + f(52) - f(13) - f(39)$$

In other words, k measures the difference in frequencies between even-cycled $m^6A$ distances and odd-cycled ones. In sample AB|BA, all $m^6A$'s are expected to occur in even cycles, and $k_{AB|BA}$ should ideally be 1. In sample AB-BA, F(13):F(26):F(39) should follow a 25:50:25 distribution. So $k_{AB-BA}$ is expected to be 0. The overall contrast is defined as:

$$K = k_{AB|BA} - k_{AB-BA}$$

such that K=1 in the case of perfect contrast between the two samples. For each benchmarked method, a modification threshold scan from $P(m^6A)=0.1$ up to 0.9 is performed to determine the best K.

### GLORI benchmark (HEK293)

dRNA-Seq reads are basecalled and then aligned to genome GRCh38. Reads with mapping quality below 50 are filtered out. Read-level $P(m^6A)$ is thresholded at 0.5 to classify each nucleotide on a molecule into modified or unmodified status. All predictions that align to the same site as listed in GLORI are then aggregated to produce the mAFiA site-level stoichiometry. To ensure values from both sides are reasonably confident, only GLORI sites with adjusted p-values less than 0.001 are included. On the other hand, a minimum dRNA-Seq coverage of 50 is required for a site to be compared. The intersection of these two requirements yields a total of 5762 sites.

The complete code and steps for reproducing the results are available at: https://github.com/dieterich-lab/mAFiA

### Isoform-specific Predictions

The transcriptome GRCh38.102 is used to assign each $m^6A$ locus on a read to a specific isoform. Only isoforms with coverage of higher than 10 are put forward for stoichiometry calculation. dRNA-Seq data is obtained from HEK293 WT in two replicates. Replicate 1 is performed on an ONT MinION, replicate 2 on ONT P2.

### Comparison to CHEUI and m6Anet

Basecalling for CHEUI and m6Anet is done with ONT Guppy 6.4.6, execpt for AB-BA design oligos, which is done with Guppy 6.5.7. The results are mapped to the reference transcriptome GRCh38.102 with minimap 2.22. Resquiggling is done with nanopolish 0.13.3. The remaining procedures follow the documentations of CHEUI and m6Anet 2.0.1. Read-level predictions are used for the single-molecule benchmark, while site-level predictions are used for HEK293. For GLORI sites that map to multiple transcripts, a weighted average of the isoforms is calculated.

The 3-way intersection of GLORI sites covered by mAFiA, CHEUI and m6Anet yields a subset of 2240 sites. Only these sites are used to compute a method’s correlation to the GLORI reference as listed in Table 4.

### Supplementary Figures

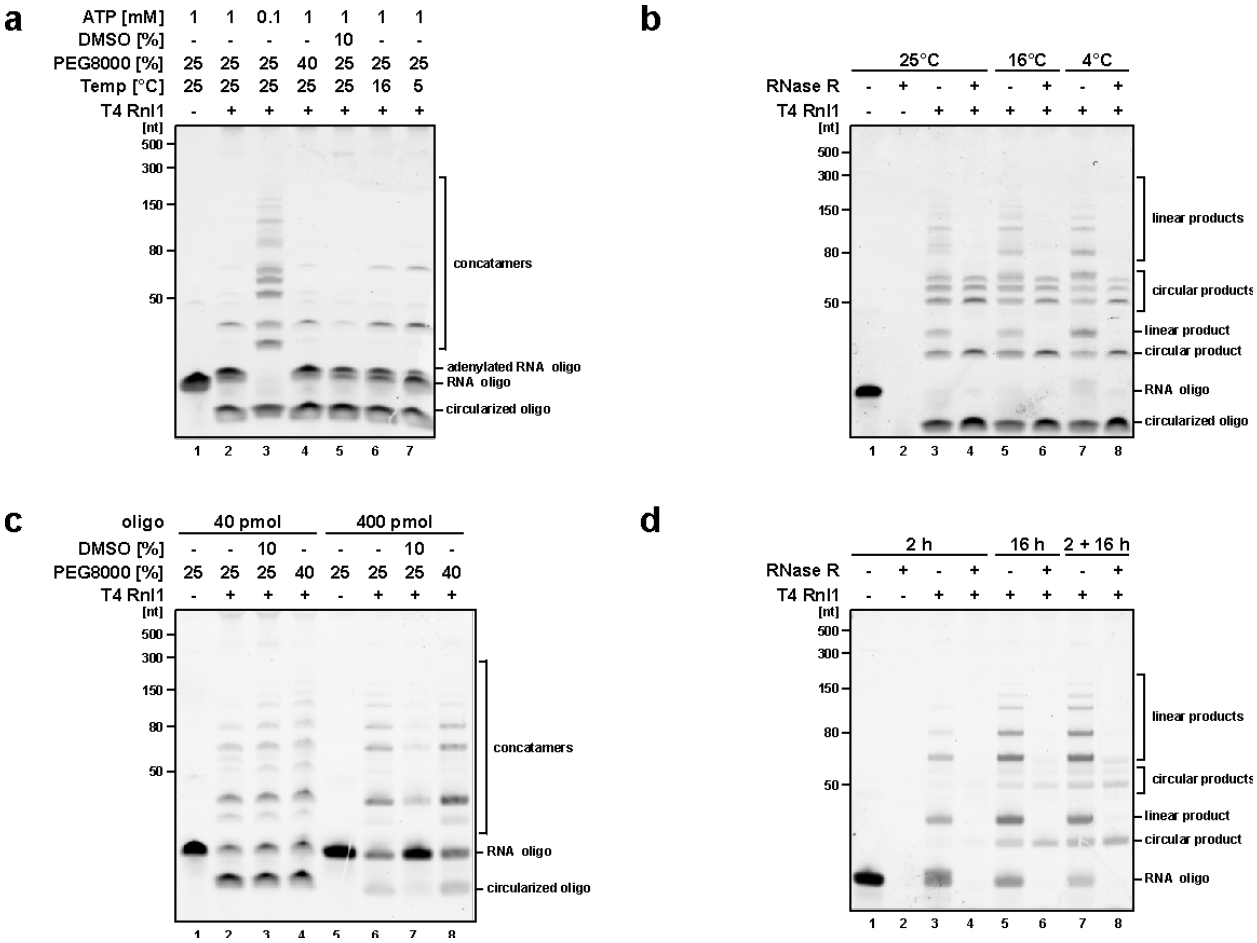


#### Figure S1: Optimization of Random Ligation approach

**a,** Influence of ATP concentration, DMSO addition, PEG8000 concentration and reaction temperature was tested for random ligation of oligo RL-M1S0. Reaction time = 2 h.

**b,** Test of different reaction temperatures for Random Ligation of RL-M1S0 as indicated. All linear reaction products were digested with RNase R where indicated. Reaction time = 2h.

**c,** Influence of the oligo concentration, DMSO addition and PEG8000 concentration on the Random Ligation. Reaction was carried out for 2 h at 4°C.

**d,** Test of different reaction times as indicated. For the 2+ 16 h approach, fresh enzyme and ATP were added after 2 h. Reactions were carried out at 4°C and afterwards digested with RNase R where indicated.

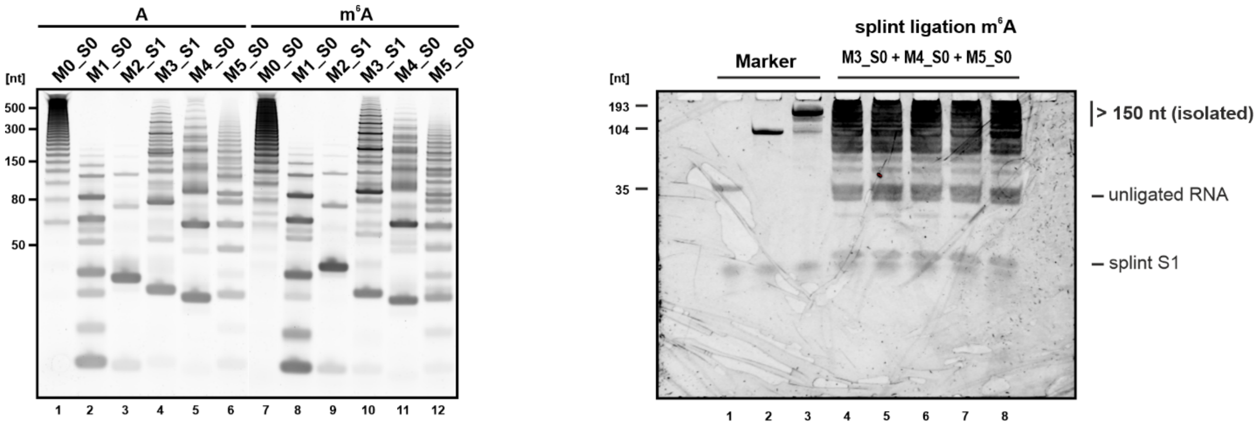


**Figure S2: Final Random Ligation (left) and Splinted Ligation (right).**

Exemplary gel images of RL and SPL sample preparation. The sequences of RNA oligos are listed in Table 1, and contain either an A or m$^6$A in the central position. Oligos were ligated according to the optimized Random Ligation protocol and analyzed by TBE-urea PAGE.

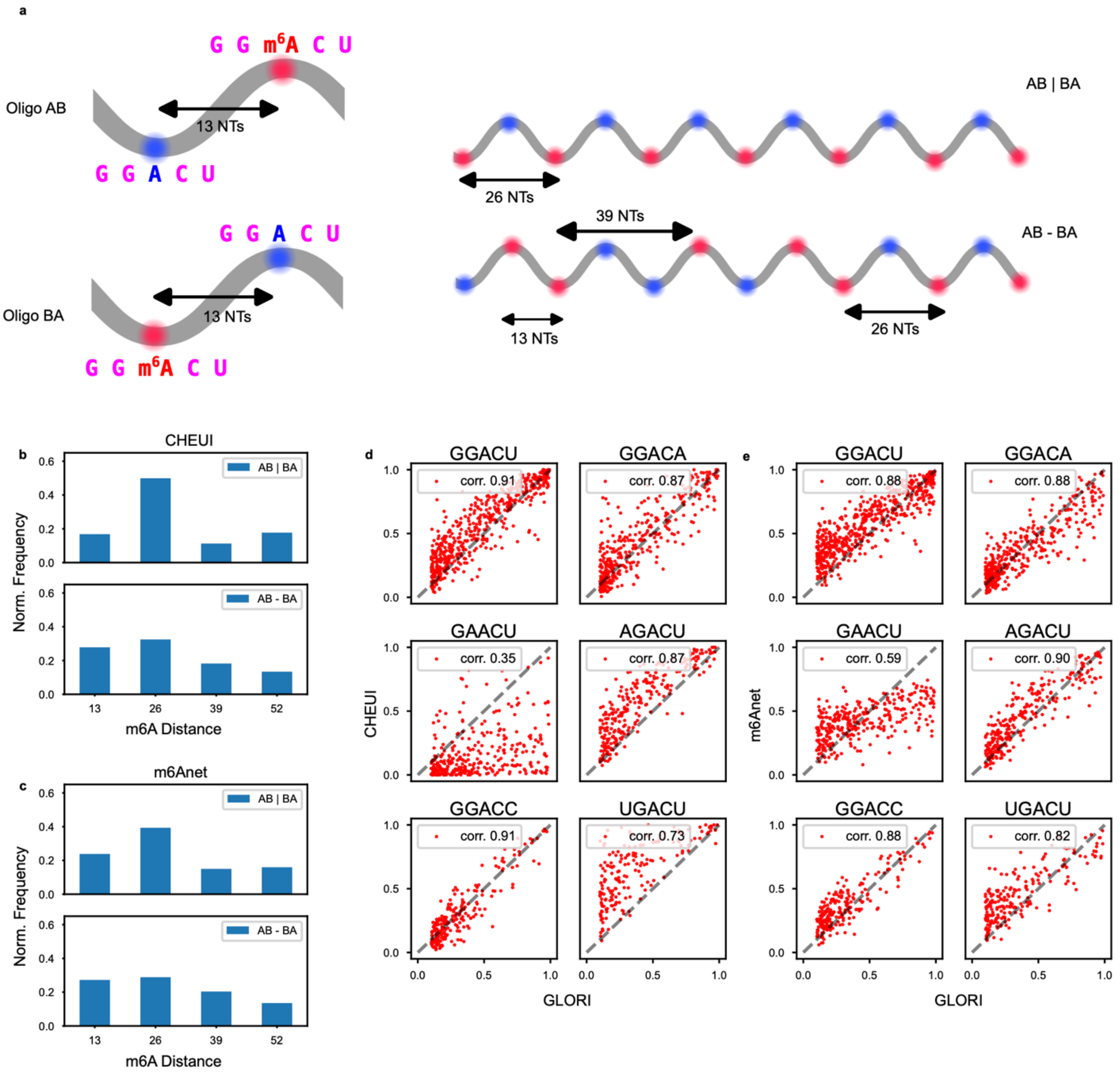


**Figure S3: Benchmarking CHEUI and m6Anet on synthetic molecule and HEK293 data.**

**a,** Illustration of the ligation products AB|BA and AB-BA, which contain 26-mer oligos of patterns AB and BA in different arrangements. A and B are identical 13-nt sequences containing the motif GGACU, except that sequence A has an unmodified A in its center, while B has an $m^6A$ (Table 2). In the first sample AB|BA, the two sets of oligos are ligated separately and then mixed together, whereas in the second sample AB-BA, both oligos are ligated into heteropolymers with arbitrary order. The former sample only allows $m^6A$ spacing of 26 nts, while the latter admits all spacing in multiples of 13 nts.

**b,** Distribution of m$^6$A distance per-read predicted by CHEUI on AB+BA, with contrast score 0.40.

**c,** Distribution of m$^6$A distance per-read predicted by m6Anet on AB+BA, with contrast score 0.22.

**d,** Site-level modification ratio on HEK293, CHEUI vs GLORI, overall correlation 0.64.

**e,** Site-level modification ratio on HEK293, m6Anet vs GLORI, overall correlation 0.80.

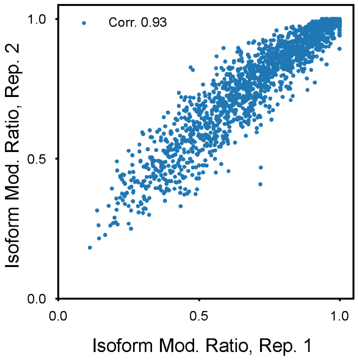


**Figure S4: Reproducibility of isoform-specific modification ratio.** Replicate 1 is obtained from ONT MinION with 1.3M reads after filtering. Replicate 2 from ONT P2 with 3.5M reads.

| | RL (21-mer) | SPL (33-mer) |
|---|---|---|
| M0_S0 | AUUGUCAUGGA/$m^6$ACUCUGGAGAC | GGCGUGAUUGUCAUGGA/$m^6$ACUCUGGAGACCAACUA |
| M1_S0 | UGCACAGAGGA/$m^6$ACAAGUAGCUG | GGCGUGUGGGAGAUGGA/$m^6$ACAAUAUGCUCCAACUA |
| M2_S0 | UACUGAAGGAA/$m^6$ACUGACCUCUC | GGCGUGUACUGAAGGAA/$m^6$ACUGACCUCUCCAACUA |
| M2_S1 | UAGUAAAGGAA/$m^6$ACUAUGCAAAU | |
| M3_S0 | GAAUGUAUAGA/$m^6$ACUUUAUGUCU | GGCGUGGUAGUAAAAGA/$m^6$ACUGGUUAAUGCAACUA |
| M3_S1 | UUCCCAAGAGA/$m^6$ACUGAGCGGGC | |
| M4_S0 | CCCCAGCUGGA/$m^6$ACCGACUCAGA | GGCGUGUGCAGUUAGGA/$m^6$ACCUCAGUGGCCAACUA |
| M5_S0 | GGGACCUCUGA/$m^6$ACUGCUCUGGG | GGCGUGGGGACCUCUGA/$m^6$ACUGCUCUGGGCAACUA |

**Table 1: List of oligo designs used for training.** Modified A highlighted in red, unmodified in blue, surrounding motif in purple.

| SPL-AB | SPL-BA |
|---|---|
| GGCAGGACUGCUAGGCAGG$m^6$ACUGCUA | GGCAGG$m^6$ACUGCUAGGCAGGACUGCUA |

**Table 2: Oligo designs used for benchmarking single-molecule $m^6$A detection.**

| splint D1 for ligation of M0_S0 to M5_S0 | splint D2 for ligation of AB and BA oligos |
|---|---|
| CACGCCTAGTTG | CCTGCCTAGCAG |

**Table 3: Splint designs for ligation of SPL molecules**

| | AB+BA | GLORI |
|---|---|---|
| mAFiA | **0.66** | **0.85** |
| CHEUI | 0.40 | 0.64 |
| m6Anet | 0.22 | 0.80 |

**Table 4: Summary of benchmark results.** mAFiA outperforms other single-read $m^6A$ detection methods in both single-molecule and mRNA benchmarks (best score for each benchmark in bold).
